# Controlled-release urea combined with potassium chloride improved the soil fertility and growth of Italian ryegrass

**DOI:** 10.1101/2020.06.13.150318

**Authors:** Jibiao Geng, Xiuyi Yang, Xianqi Huo, Jianqiu Chen, Shutong Lei, Hui Li, Ying Lang, Qianjin Liu

**Affiliations:** Shandong Provincial Key Laboratory of Water and Soil Conservation and Environmental Protection, State Key Laboratory of Nutrition Resources Integrated Utilization, College of Agriculture and Forestry Science/Resources and Environment, Linyi University, Linyi, Shandong 276000, China; Kingenta Ecological Engineering Group Co., Ltd., Linshu, Shandong 276700, China

**Keywords:** Controlled-release urea, growth characteristics, Italian ryegrass, potassium chloride, soil fertility, yield

## Abstract

A field experiment with a split-plot design was conducted to study the effect of nitrogen fertilizer type combined with different potassium fertilizer rates on the soil fertility and growth of Italian ryegrass. The main plots were assigned to controlled-release urea (CRU) and common urea, while low, moderate and high potassium chloride (KCl) rates (150, 300 and 450 kg ha^−1^, respectively) were assigned to the subplots. The results showed compared with the common urea, the CRU significantly increased the SPAD value, plant height, leaf area, and photosynthetic index. Moreover, the dry and fresh yields of the CRU increased by 10.9-25.3% and 11.8-17.7%, respectively. At the same time, compared with the KCl150 and KCl450 treatments, the KCl300 treatment resulted in better plant growth. Overall, the CRU×KCl300 maximized the soil inorganic nitrogen and different soil potassium forms. The root length, volume, surface area, average diameter, tips and branches were also improved, and there was a significant N×K interaction effect on the tips. Our analysis corroborated the CRU combined with 300 kg ha^−1^ KCl fertilization enhances crop growth by improving leaf photosynthesis, soil fertility, and yield and should be recommended as the best fertilizer ratio for Italian ryegrass production.

## Introduction

Grassland accounts for 41.7% of China’s total land area (Liu *et al*., 2018). For a long time, as the main means of production and material basis, grasslands have made important contributions to the development of animal husbandry, and grasslands have an important ecological function that plays a central role in water and soil conservation, air purification, climate regulation and biodiversity conservation (Binder *et al*., 2018). Italian ryegrass (*Lolium multiflorum* L.) is a globally cultivated grass. This grass has many tillers at its roots, grows quickly and has good grazing resistance. Italian ryegrass has a strong adaptability and high yield and nutrition, which play a role in improving soil (Hussain *et al*., 2018; Svatos and Abbott, 2019). Italian ryegrass is widely used in parks, golf courses and other recreational sporting areas as the first choice of lawn grass in foreign countries (Fan *et al*., 2018). Various production technologies and late management measures have also become new research topics. At present, research on ryegrass in lawns, playgrounds and other fields in China is immature. A large number of researchers have focused on management technology after ryegrass planting (Alves dos Santos *et al*., 2018; He *et al*., 2020).

Nitrogen is not only the basis of forage genetic material but also the composition of many important organic compounds (Bolinder *et al*., 2010). It is very important for life cycle activities and the yield and nutritional quality of forage. Italian ryegrass does not have the ability to fix nitrogen, so nitrogen in the soil is a key factor in grass growth and development (Woods *et al*., 2018). The application of nitrogen fertilizer can significantly improve the yield of Italian ryegrass, and the plant can absorb both ammonium and nitrate nitrogen (Cavalli *et al*., 2016). Moreover, Italian ryegrass is a fast-growing forage grass with a high N requirement and, therefore, strongly relies on N soil content to maintain adequate forage yield (Masoni *et al*., 2015).

The fertilizer utilization rate in China is generally low, mainly due to the rapid dissolution of instant fertilizer after it is applied to the soil (Wang *et al*., 2018). The crop cannot absorb and utilize nitrogen in time, which results in the loss of most of the nitrogen in gaseous or water-soluble forms and causes a series of environmental problems, such as eutrophication of surface water, nitrate pollution of groundwater and agricultural products, and ammonia and nitrogen oxides emitted to the ozone layer (Ata-Ul-Karim *et al*., 2017). In addition, it is difficult to apply fertilizer to Italian ryegrass after every cutting, so decreasing the fertilizer application frequency and improving the yield and quality of Italian ryegrass are the key problems to address in the planting process (Martin *et al*., 2017). An effective solution may be to develop a new type of slow-release fertilizer to meet the needs of crop growth.

In recent years, controlled-release urea (CRU) has been used worldwide (Geng *et al*., 2016; Li *et al*., 2018; Liu *et al*., 2019). CRU can release nitrogen slowly in the form of a resin polymer coating, continuously supply nutrients needed for ryegrass growth, decrease the number of topdressings and labour intensity in the later stage, simplify cultivation technology, save time and labour, and reduce environmental pollution (Gaylord *et al*., 1975).

Potassium chloride (KCl) is commonly used in agricultural production. It has a low price and high nutrient content, and the application of appropriate amounts can promote the growth of Italian ryegrass (McDonnell, *et al*., 2018). In addition, potassium can improve the photosynthetic capacity and disease resistance of plants and then extend the green period (Hasanuzzaman *et al*., 2018). In addition, a single application of nitrogen and potassium or mixed application of different proportions can increase the fresh yield of Italian ryegrass, but the effect of a mixed application is better than that of a single application (Oliveira *et al*., 2017). According to the supply and demand curve of soil for nutrient elements, the most appropriate fertilization amount and fertilization type can be formulated.

Nitrogen can improve photosynthesis and thus dry matter production (Jarvis, 1987). Potassium is very important for root growth and disease resistance (Snyder and Cisar, 2000). The possible interactions between the two nutrients are unknown, as they have not been sufficiently studied in Italian ryegrass. In previous studies on the effect of N×K fertilizer application on crop growth, the positive interaction of N and K reduced the cost of fertilizer and contributed to food security (Yang *et al*., 2016a, 2016b). Nitrogen fertilizer application increases the plant potassium absorption efficiency, and potassium fertilizer application resolves the problem of nitrogen pollution by inducing in crops a high nitrogen absorption efficiency (Dong *et al*., 2010). Eventually, the mutual promotion of nitrogen and potassium enhances the yield and quality of crops (Yang *et al*., 2016a). This is a feasible way to increase the potassium fertilizer input and improve the nitrogen utilization efficiency. Understanding the mechanism underlying the N×K interaction is vital to guide the best practice of nutrient management in agricultural production.

Furthermore, there are few reports on the interaction application effects of CRU combined with KCl on Italian ryegrass growth. Hence, the objective of this study was to investigate the effects of CRU in combination with KCl on (i) soil inorganic nitrogen (NO_3_^−^-N and NH_4_^+^-N), (ii) soil potassium forms (available potassium, water-soluble potassium, exchangeable potassium and nonexchangeable potassium), (iii) growth and photosynthesis characteristics, and (iv) Italian ryegrass yield and fertilizer use efficiency.

## Materials and methods

### Experimental site and material

The field experiment was arranged at the experimental base of the College of Agriculture and Forestry Science, Linyi University, Linyi city, Shandong Province (35°06’N, 118°17’E) in 2019, and this site has a continental climate typical of temperate monsoon areas. The temperature and relative humidity were 30 ± 5 °C and 45 ± 5% mm, respectively. The experimental site has four distinct seasons and substantial light. The tested soil is sandy loam, which is classified as Typic Hapludalf according to the USDA classification (Soil Survey Staff 1999). The basic soil properties are listed in Table 1.

**Table 1.** Part properties of tested soil before Italian ryegrass planting in 2019.

| pH value (2.5:1) | Organic matter ($\text{g kg}^{-1}$) | Total N ($\text{g kg}^{-1}$) | $\text{NO}_3^-$-N ($\text{mg kg}^{-1}$) | $\text{NH}_4^+$-N ($\text{mg kg}^{-1}$) | Available P ($\text{mg kg}^{-1}$) | Available K ($\text{mg kg}^{-1}$) |
| --- | --- | --- | --- | --- | --- | --- |
| 6.52 | 6.88 | 0.73 | 46.09 | 31.22 | 39.82 | 182.08 |

The tested forage is ‘Bluesign’ Italian ryegrass, which is produced by Suqian Chengzhiyang Seed Industry Co., Ltd., China. The seeding rate of the Italian ryegrass was 25 kg ha^−1^. The fertilizers used included CRU and ordinary fertilizers. The CRU (containing N 43%, with a release period of almost 3 months in distilled water at 25 °C) (Fig. 1) fertilizer was produced by the State Key Laboratory of Nutrition Resources Integrated Utilization, China. The CRU fertilizer was formulated as round particles with a regular shape and smooth surface, and it was coated with an epoxy resin (low curing shrinkage, strong adhesion and chemical resistance). The ordinary fertilizers included urea (containing N 46%), calcium superphosphate (containing P_2_O_5_ 14%), and KCl (containing K_2_O 60%), which were provided by Jinzhengda Ecological Engineering Group Co., Ltd., and Kingenta Ecological Engineering Group Co., Ltd., China.

**Fig. 1.**
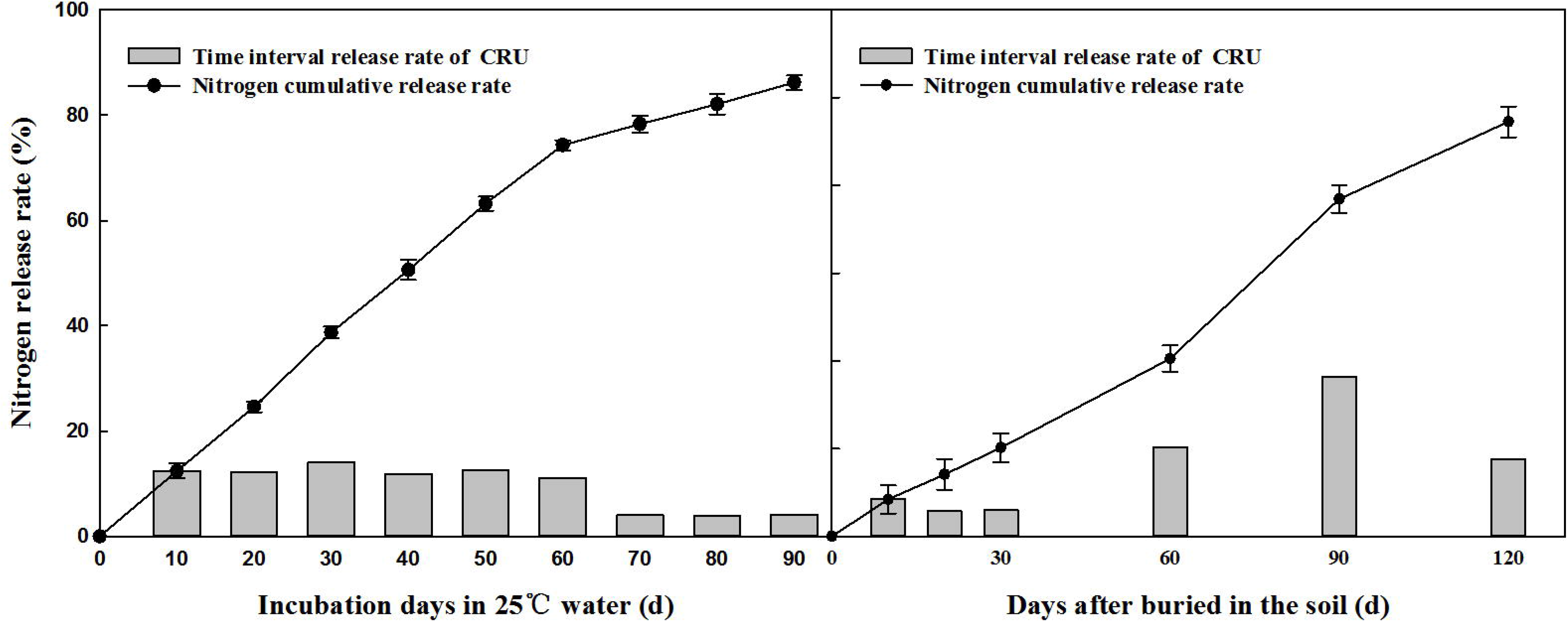
Release of nitrogen from CRU.

### Experimental design

A split-plot design with three replications was used for the experiment. Specifically, the nitrogen types (CRU and urea) were the main plots, KCl rates (150, 300 and 450 kg ha^−1^) were the subplots, and no N or K fertilization was the control. The main plot covered an area of 45 m^2^ (3 m wide and 15 m long), and the subplot covered an area of 15 m^2^ (3 m wide and 5 m long). Each plot received a basal application of 300 kg ha^−1^ N and 200 kg ha^−1^ P_2_O_5._ All fertilizers (except common urea) were applied once with hands before sowing seeds. In particular, urea was applied twice in total, with 60% before sowing seeds and 40% after the second clipping stage. The Italian ryegrass was sown on April 4, 2019. CRU (10 g) fertilizer was weighed and placed into the mesh bag (8 cm width and 10 cm length), the bags were sealed, and this process was repeated 18 times. Then, these bags were put into the ploughed soil layer before the Italian ryegrass was sown to determine the release characteristics of the CRU fertilizer buried in the soil.

### Sampling and measurement

The release of the CRU in the soil was determined by the buried bag method. Similarly, 10 g of CRU particles were put in a mesh bag (8 cm wide and 10 cm long) buried in a cement tank at a depth of 15-20 cm during fertilization. The bags were collected on the 10^th^, 20^th^, 30^th^, 60^th^, 90^th^, and 120^th^ days after burial. Three bags were collected each time, washed and dried to constant weight at 60 °C, and the nitrogen release rate was calculated according to the weight of the remaining fertilizer particles.

Soil and plant samples were collected on May 18, 2019 (first clipping), June 14, 2019 (second clipping), July 16, 2019 (third clipping), and August 10, 2019 (fourth clipping), and the physiological indexes of Italian ryegrass under different treatments were observed and measured.

The 0-20 cm soil samples were collected by a 5-point sampling method (2 sampling points in the fertilizer row, 3 sampling points in the plant row). The contents of NO_3_^−^-N and NH_4_^+^-N in fresh soil (0.01 mol L^−1^ CaCl_2_ extraction) were determined immediately using an AA3 continuous flow analyser (Bran-Luebbe, Norderstedt, Germany). The remaining soil was dried by naturally existing air and ground through 2 mm and 0.25 mm sieves, and the organic matter (potassium dichromate external heating method), soil total N (semi micro Kelvin method), available phosphorus (pH 8.5, 0.5 mol L^−1^ NaHCO_3_ extraction, molybdenum blue colorimetry) and available potassium (1 mol L^−1^ NH_4_OAc extraction, flame photometer method) contents were determined (Zheng *et al*., 2016).

The leaf area of Italian ryegrass was measured by a leaf area meter (Yaxin-1241, Yaxinliyi, China). The SPAD value was measured by a hand-held chlorophyll meter (SPAD-502, Minolta, Japan). Besides, the Li-6400 portable photosynthetic apparatus (LI-COR, Lincoln, NE, USA) was also used for the determination. The leaf photosynthetic indicators were measured from 9:00-10:00 a.m. Under sunny and cloudless weather, the net photosynthetic rate (*P*_n_), stomatal conductance (*G*_s_), intercellular carbon dioxide concentration (*C*_*i*_) and transpiration rate (*T*_r_) were measured before the second clipping.

After measuring the physiological indexes of the plant, ten successive plants above the roots from each subplot were clipped with scissors, and the height of the stubble was the same as that of the ground. The fresh weight of the yield of each treatment was recorded. Then, the clipped grass was sealed in the file bag according to the treatment label, placed in the oven at 105 °C for 30 minutes, and dried at 65 °C for 72 hours, and then, the dry weight of the yield was recorded. Finally, the nitrogen and potassium contents of the plants were analysed. The plant total nitrogen contents were determined by digestion with H_2_SO_4_-H_2_O using the micro-Kjeldahl method. The potassium content of the plant was digested by H_2_SO_4_-H_2_O and determined by a flame photometer. The nitrogen and potassium uptake were calculated according to the nitrogen and potassium content and dry mass weight of each plot. The nitrogen use efficiency (NUE) and potassium use efficiency (KUE) were calculated (Rietra *et al*., 2017).

The root samples were scanned with a flatbed image scanner (Epson Expression/STD LC-4800 scanner). The images were analysed by WinRHIZO commercial software (Regent Instruments, 2001) to determine the root volume, total length, diameter, surface area, and numbers of tips and branches.

### Statistical analyses

Microsoft Excel 2010 was employed for data processing, and Sigma Plot software version 10 (MMIV, Systat Software Inc., San Jose, CA, USA) was used to draw the figures. Data were subjected to analysis of variance (ANOVA) and mean separation tests as a split-plot factorial design with three replications. Concretely, the data were analysed using Statistical Analysis System version 9.2 (SAS Institute Cary, NC, 2010) with a two-way ANOVA at a significance level of 0.05, with nitrogen type and potassium rate as the independent variables. Two-way ANOVAs were performed to determine the effects of N, K and their interactions on the leaf area, leaf SPAD, leaf photosynthesis chlorophyll parameters, root morphology, yield and nutrient uptake of Italian ryegrass. One-way ANOVAs were performed to test for significant differences between treatments of NH_4_^+^-N, NO_3_^−^-N, and soil K informs. A Duncan multiple range test was carried out to determine if significant (p<0.05) differences occurred between individual treatments (Tang and Feng, 2002).

## Results

### Release characteristics of the CRU fertilizer

The release characteristics of the CRU fertilizer in the soil were released in the form of “S”, reaching 80% in approximately 100 days (Fig. 1). The release period of the CRU in the soil was almost 120 days: the release rate was slow within 0~60 days, the release rate increased within 60~90 days, and the nutrient decline period was within 90~120 days, which met the nitrogen demand of Italian ryegrass.

### Soil inorganic nitrogen and potassium form

The contents of NO_3_^−^-N and NH_4_^+^-N in the soil in the control treatment were the lowest among all the treatments in the whole plant growing season (Fig. 2). With the increase in the KCl application rate, the contents of NO_3_^−^-N and NH_4_^+^-N in the soil changed little, which was not related to the type of nitrogen application. However, at the early stage of growth, the contents of NO_3_^−^-N and NH_4_^+^-N in the urea treatment were higher than those in the CRU treatment, but after the second clipping, the contents of NO_3_^−^-N and NH_4_^+^-N in the urea treatment decreased rapidly; in addition, the contents of NO_3_^−^-N and NH_4_^+^-N in the urea treatment were lower than those in the CRU treatment.

**Fig. 2.**
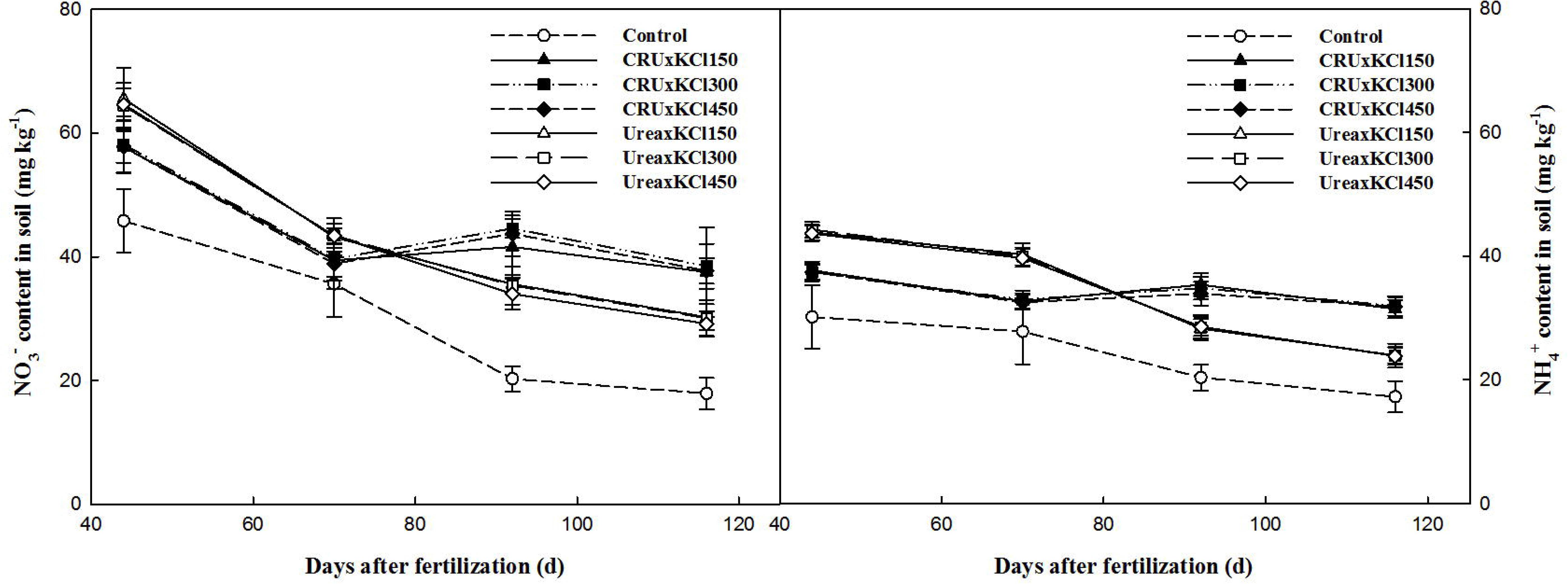
Change of the soil NO_3_^−^-N and NH_4_^+^-N contents.

Overall, the contents of available K, water-soluble K, exchangeable K and nonexchangeable K were significantly affected by the KCl application rate, and of the treatments, the control treatment had the lowest values in different developmental stages (Fig. 3). The contents of soil available K, water-soluble K and nonexchangeable K decreased gradually, but that of exchangeable K showed a fluctuating trend. Regardless of which type of nitrogen fertilizer was combined, the contents of soil available potassium, water soluble potassium and exchangeable potassium improved with the increase in the KCl application rate, and there was a significant difference between the different potassium fertilizer treatments.

**Fig. 3.**
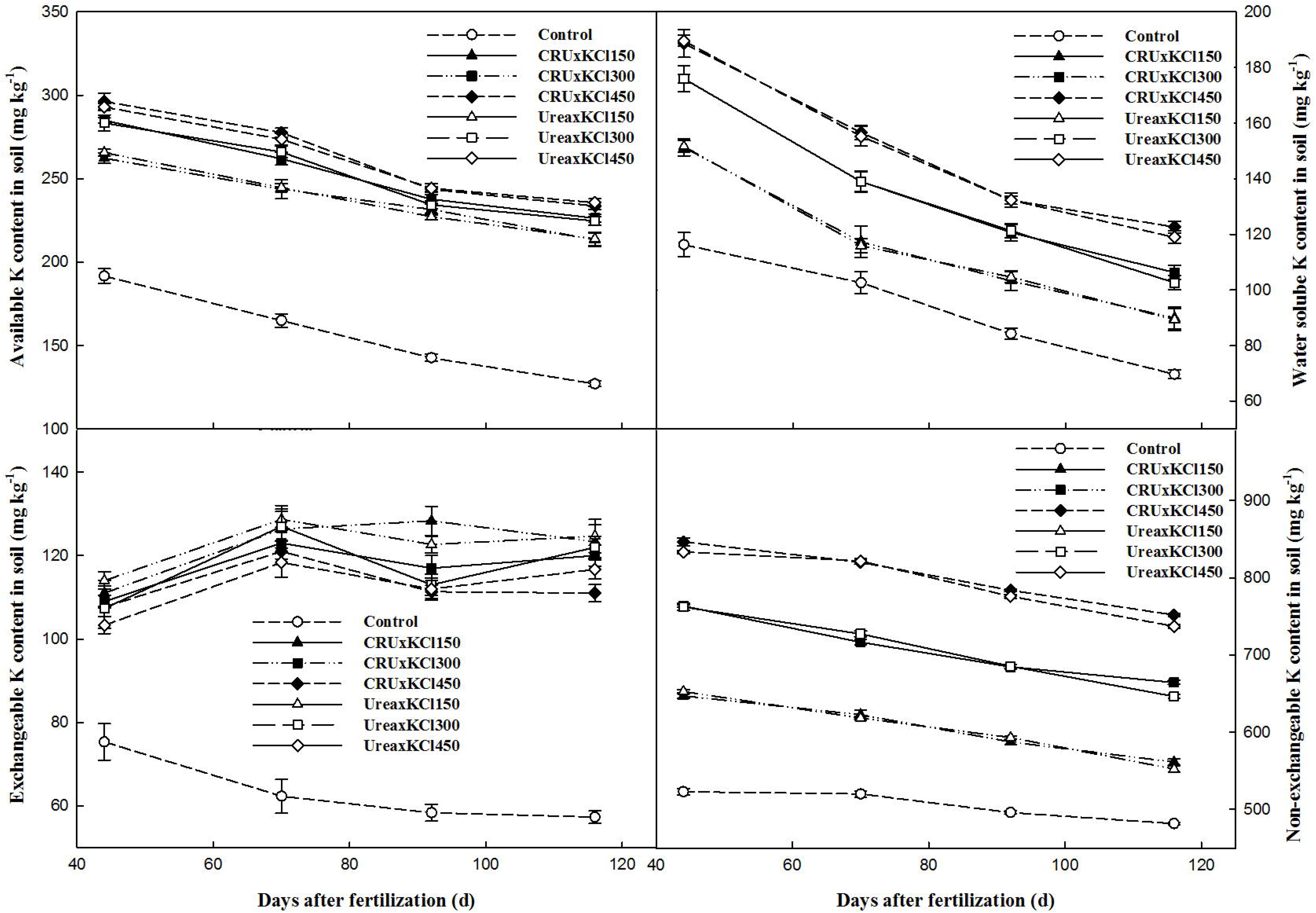
Change of soil available K, water soluble K, exchangeable K and non-exchangeable K contents.

### Plant height and leaf area

In the whole growth period of Italian ryegrass, the plant heights of the different fertilization treatments increased first and then decreased (Table 2). There was no significant difference between the different fertilization treatments at the first and second clipping stages, but the advantage of the CRU treatment group was obvious; in addition, the average value was higher than that of the control treatment. During the mowing period of the third and fourth clippings, Italian ryegrass growth was in the transition period from maturity to senescence, the plant demand for nutrients decreased, and the ability to absorb fertilizer also decreased. Under the same amount of applied nitrogen, the plants in the KCl150 and KCl300 treatments were taller, and those in the KCl450 treatment were shorter. At the same time, the plant height of the CRU treatment group was significantly greater than that of the urea treatment groups.

**Table 2.** The plant height of Italian ryegrass under different treatments.

| Treatment | First clipping (cm) | Second clipping (cm) | Third clipping (cm) | Fourth clipping (cm) |
| --- | --- | --- | --- | --- |
| **Nitrogen fertilizer type** |  |  |  |  |
| CRU | 17.64 a | 25.22 a | 24.47 a | 15.50 a |
| Urea | 16.67 a | 23.94 a | 19.83 b | 8.06 b |
| **Potassium fertilizer rate ($\text{kg ha}^{-1}$)** |  |  |  |  |
| 150 | 17.78 a | 23.47 a | 22.70 a | 11.17 b |
| **300** | **16.48 a** | **25.17 a** | **21.62 a** | **13.08 a** |
| **450** | **17.20 a** | **25.12 a** | **20.63 a** | **11.08 b** |
| **N×K interaction** |  |  |  |  |
| **Control** | **16.27 a** | **21.43 c** | **20.30 b** | **8.32 b** |
| **CRU×KCl150** | **17.83 a** | **24.57 abc** | **25.23 a** | **14.33 a** |
| **CRU×KCl300** | **16.77 a** | **25.90 a** | **23.23 ab** | **17.02 a** |
| **CRU×KCl450** | **18.33 a** | **25.20 ab** | **21.93 ab** | **15.16 a** |
| **Urea×KCl150** | **17.73 a** | **22.36 bc** | **20.17 b** | **8.32 b** |
| **Urea×KCl300** | **16.20 a** | **24.43 abc** | **20.00 b** | **9.16 b** |
| **Urea×KCl450** | **16.07 a** | **25.03 ab** | **19.33 b** | **7.03 b** |
| **Source of variance** |  |  |  |  |
| **N** | **0.1619** | **0.1735** | **0.0023** | **<0.0001** |
| **K** | **0.3002** | **0.2415** | **0.1858** | **0.0632** |
| **N×K** | **0.3855** | **0.6337** | **0.4809** | **0.5067** |

**Note:** Means followed by different lowercase letters in the same column were significantly different based on analyses with ANOVAs followed by Duncan tests ( $P < 0.05$ ).
**Control:** no nitrogen and potassium fertilizer, **CRU:** polymer-coated urea, **KCl:** potassium chloride.

Similarly, the leaf area of Italian ryegrass in the whole growth period increased first and then decreased with the growth of the plant (Table 3). At the second clipping period, the apparent leaf area increased the fastest. The leaves of the plants were long and thin, and there was no difference between the fertilization treatments at the first and second clipping stages, but fertilization was more suitable for its growth; in addition, the leaf area of the plants with fertilization was relatively large. During the third and fourth clipping stages, in comparison with the control and common urea treatments, the CRU treatments resulted in a significant difference, which indicated that the CRU could delay plant senescence to some extent. In addition, the effect of nitrogen fertilizer on ryegrass leaf area was greater than that of KCl fertilizer, and the sustainability of combined application is more important. There was no significant N×K interaction effect on the plant height and leaf area (except at the third clipping).

**Table 3.**
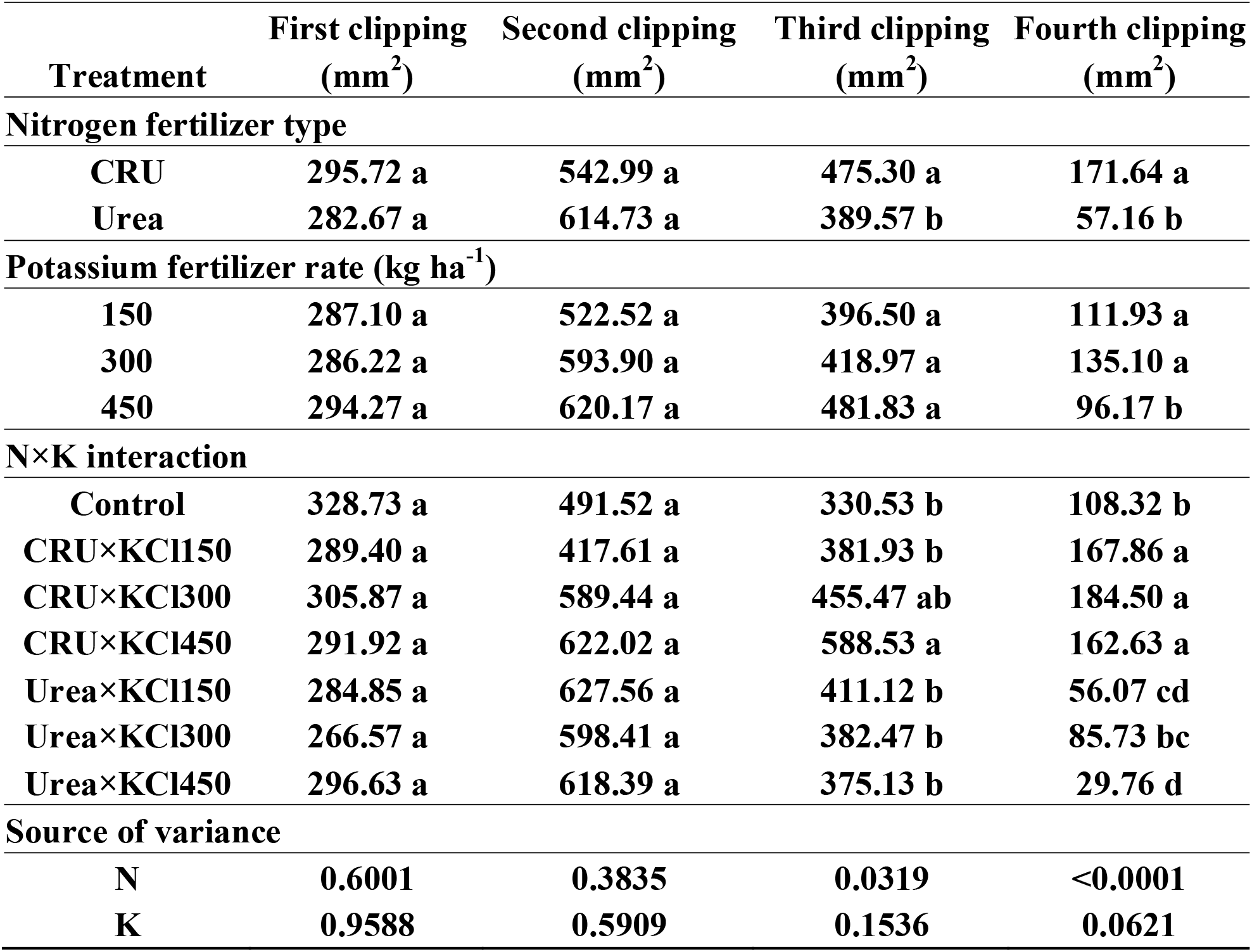
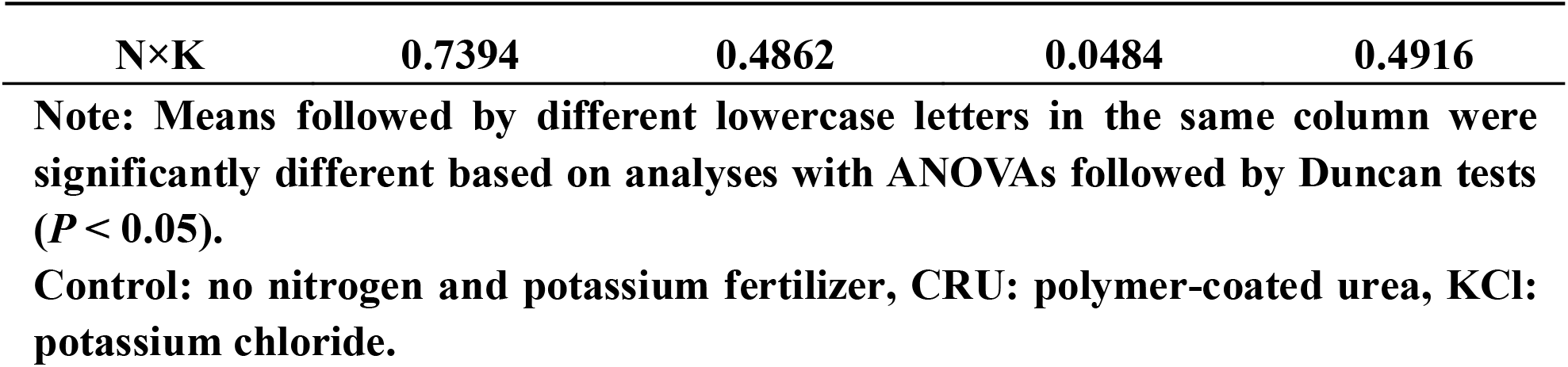
The leaf area of Italian ryegrass under different treatments.

| **Treatment** | **First clipping**<br>**(mm<sup>2</sup>)** | **Second clipping**<br>**(mm<sup>2</sup>)** | **Third clipping**<br>**(mm<sup>2</sup>)** | **Fourth clipping**<br>**(mm<sup>2</sup>)** |
| --- | --- | --- | --- | --- |
| **Nitrogen fertilizer type** |  |  |  |  |
| **CRU** | **295.72 a** | **542.99 a** | **475.30 a** | **171.64 a** |
| **Urea** | **282.67 a** | **614.73 a** | **389.57 b** | **57.16 b** |
| **Potassium fertilizer rate (kg ha<sup>-1</sup>)** |  |  |  |  |
| **150** | **287.10 a** | **522.52 a** | **396.50 a** | **111.93 a** |
| **300** | **286.22 a** | **593.90 a** | **418.97 a** | **135.10 a** |
| **450** | **294.27 a** | **620.17 a** | **481.83 a** | **96.17 b** |
| **N×K interaction** |  |  |  |  |
| **Control** | **328.73 a** | **491.52 a** | **330.53 b** | **108.32 b** |
| **CRU×KCl150** | **289.40 a** | **417.61 a** | **381.93 b** | **167.86 a** |
| **CRU×KCl300** | **305.87 a** | **589.44 a** | **455.47 ab** | **184.50 a** |
| **CRU×KCl450** | **291.92 a** | **622.02 a** | **588.53 a** | **162.63 a** |
| **Urea×KCl150** | **284.85 a** | **627.56 a** | **411.12 b** | **56.07 cd** |
| **Urea×KCl300** | **266.57 a** | **598.41 a** | **382.47 b** | **85.73 bc** |
| **Urea×KCl450** | **296.63 a** | **618.39 a** | **375.13 b** | **29.76 d** |
| **Source of variance** |  |  |  |  |
| **N** | **0.6001** | **0.3835** | **0.0319** | **<0.0001** |
| **K** | **0.9588** | **0.5909** | **0.1536** | **0.0621** |

| N×K | 0.7394 | 0.4862 | 0.0484 | 0.4916 |
| --- | --- | --- | --- | --- |

Note: Means followed by different lowercase letters in the same column were significantly different based on analyses with ANOVAs followed by Duncan tests ( $P < 0.05$ ).

### Leaf SPAD and photosynthetic index

The effect of different fertilization treatments on the SPAD value of Italian ryegrass was different (Table 4). At the first clipping stage, the SPAD value of the CRU×KCl300 treatment was the highest, 17.1% higher than that of the control treatment. During the whole growth period of Italian ryegrass, the SPAD value increased first and then decreased, and the SPAD value of the CRU treatment was higher than that of the common urea treatment. Under the three levels of KCl fertilizer, the SPAD value of the KCl150 treatment in the whole growth period was low, which indicates that the insufficient use of potassium had a certain impact on the SPAD value. The SPAD value of the CRU×KCl300 treatment was the highest at the third and fourth stages, while the effect of nitrogen on the SPAD value of ryegrass was greater at the later stage. In comparison with common urea, the application of CRU had a greater advantage in improving the SPAD value. There was no significant N×K interaction effect on the SPAD value (except at the second clipping).

**Table 4.** The leaf SPAD value of Italian ryegrass under different treatments.

| Treatment | First clipping | Second clipping | Third clipping | Fourth clipping |
| --- | --- | --- | --- | --- |
| Nitrogen fertilizer type |  |  |  |  |
| CRU | 49.76 a | 54.70 a | 27.49 a | 17.44 a |
| Urea | 44.37 b | 57.66 b | 20.92 b | 10.39 b |
| Potassium fertilizer rate (kg ha<sup>-1</sup>) |  |  |  |  |
| 150 | 45.72 b | 54.83 a | 24.17 a | 14.33 a |
| 300 | 46.47 ab | 57.73 a | 25.97 a | 15.02 a |
| 450 | 49.00 a | 55.97 a | 22.48 a | 12.40 a |
| N×K interaction |  |  |  |  |
| Control | 45.67 c | 50.93 c | 25.97 ab | 13.66 b |
| CRU×KCl150 | 50.00 ab | 51.66 c | 27.56 ab | 14.86 ab |
| CRU×KCl300 | 47.66 bc | 59.62 a | 29.70 a | 22.43 a |
| CRU×KCl450 | 51.60 a | 52.74 bc | 25.20 ab | 15.03 ab |
| Urea×KCl150 | 41.43 d | 58.02 ab | 20.76 ab | 13.80 b |
| Urea×KCl300 | 45.27 c | 55.80 abc | 22.23 ab | 7.60 b |
| Urea×KCl450 | 46.40 bc | 59.16 a | 19.76 b | 9.77 b |
| Source of variance |  |  |  |  |
| N | 0.0008 | 0.0956 | 0.002 | 0.0047 |
| K | 0.0746 | 0.3603 | 0.2125 | 0.507 |
| N×K | 0.1102 | 0.0436 | 0.8487 | 0.039 |

Note: Means followed by different lowercase letters in the same column were significantly different based on analyses with ANOVAs followed by Duncan tests ( $P < 0.05$ ).
Control: no nitrogen and potassium fertilizer, CRU: polymer-coated urea, KCl: potassium chloride.

The nitrogen fertilizer types and KCl rates affected the photosynthesis indicators, but there was no significant difference between their interaction effects (Table 5). In comparison to the urea treatments, the CRU treatment improved the *P*_n_, *G*_s_ and *T*_r_ and lowered the *C*_i_. In addition, the photosynthesis indicators increased with increasing KCl rates. There was no significant N×K interaction effect on the photosynthesis indicators (except *T*_r_), and of the treatments, the CRU×KCl300 treatment performed the best in terms of Italian ryegrass leaf photosynthesis.

**Table 5.** The leaf photosynthesis chlorophyll parameters of Italian ryegrass under different treatments at the second clipping stage.

| | $P_n$ | $G_s$ | $T_r$ | $C_i$ |
| --- | --- | --- | --- | --- |
| Treatment | ($\mu\text{mol m}^{-2} \text{s}^{-1}$) | ($\mu\text{mol mol}^{-1}$) | ($\mu\text{mol m}^{-2} \text{s}^{-1}$) | ($\mu\text{mol m}^{-2} \text{s}^{-1}$) |
| Nitrogen fertilizer type |  |  |  |  |
| CRU | 295.72 a | 542.99 a | 475.30 a | 171.64 a |
| Urea | 282.67 a | 614.73 a | 389.57 b | 57.16 b |
| **Potassium fertilizer rate (kg ha<sup>-1</sup>)** |  |  |  |  |
| 150 | 287.10 a | 522.52 a | 396.50 a | 111.93 a |
| 300 | 286.22 a | 593.90 a | 418.97 a | 135.10 a |
| 450 | 294.27 a | 620.17 a | 481.83 a | 96.17 b |
| **N×K interaction** |  |  |  |  |
| Control | 328.73 a | 491.52 a | 330.53 b | 108.32 b |
| CRU×KCl150 | 289.40 a | 417.61 a | 381.93 b | 167.86 a |
| CRU×KCl300 | 305.87 a | 589.44 a | 455.47 ab | 184.50 a |
| CRU×KCl450 | 291.92 a | 622.02 a | 588.53 a | 162.63 a |
| Urea×KCl150 | 284.85 a | 627.56 a | 411.12 b | 56.07 cd |
| Urea×KCl300 | 266.57 a | 598.41 a | 382.47 b | 85.73 bc |
| Urea×KCl450 | 296.63 a | 618.39 a | 375.13 b | 29.76 d |
| **Source of variance** |  |  |  |  |
| N | 0.6001 | 0.3835 | 0.0319 | <0.0001 |
| K | 0.9588 | 0.5909 | 0.1536 | 0.0621 |
| N×K | 0.7394 | 0.4862 | 0.0484 | 0.4916 |

Note: Means followed by different lowercase letters in the same column were significantly different based on analyses with ANOVAs followed by Duncan tests ( $P < 0.05$ ).
Control: no nitrogen and potassium fertilizer, CRU: polymer-coated urea, KCl: potassium chloride, $P_n$ : photosynthetic parameters including net photosynthetic rate, $G_s$ : stomatal conductance, $C_i$ : intercellular carbon dioxide concentration, $T_r$ : transpiration rate.

### Root morphology

The nitrogen fertilizer types and potassium fertilizer rates significantly affected the root morphology (total length, surface area, average diameter, root volume, and numbers of tips and branches) (Table 6). Concretely, the CRU treatments increased these parameters compared with the urea treatments. In addition, the moderate KCl rate treatments markedly improved the root morphology compared with the low- and high-KCl-rate treatments despite any nitrogen fertilizer type. However, there was no significant N×K interaction effect (except the tips), and of the treatments, the CRU×KCl300 treatment performed the best in terms of Italian ryegrass root growth.

**Table 6.** The root morphology of Italian ryegrass under different treatments at the fourth clipping stage.

| Treatments | Total length<br>(cm) | Surface<br>area<br>(cm<sup>2</sup>) | Average<br>diameter<br>(mm) | Root<br>volume<br>(cm<sup>3</sup>) | Tips | Branches |
| --- | --- | --- | --- | --- | --- | --- |
| **Nitrogen fertilizer type** |  |  |  |  |  |  |
| CRU | 1480.89 a | 180.23 a | 0.35 a | 1.61 a | 14956.12 a | 16076.37 a |
| Urea | 1377.67 b | 159.56 b | 0.32 b | 1.49 b | 13947.33 b | 14872.35 b |
| **Potassium fertilizer rate (kg ha<sup>-1</sup>)** |  |  |  |  |  |  |
| 150 | 1393.00 c | 164.50 b | 0.32 b | 1.51 c | 14113.50 c | 14989.31 c |
| 300 | 1479.67 a | 176.17 a | 0.35 a | 1.60 a | 14754.17 a | 15905.04 a |
| 450 | 1415.16 b | 168.67 b | 0.33 b | 1.55 b | 14487.33 b | 15528.78 b |
| **N×K interaction** |  |  |  |  |  |  |
| Control | 1229.67 e | 75.33 e | 0.23 c | 0.90 d | 8142.02 e | 10214.56 e |
| CRU×KCl150 | 1434.09 bc | 173.66 b | 0.33 b | 1.56 bc | 14521.72 bc | 15490.34 c |
| CRU×KCl300 | 1545.33 a | 188.45 a | 0.36 a | 1.67 a | 15411.23 a | 16699.76 a |
| CRU×KCl450 | 1463.73 b | 178.76 b | 0.34 ab | 1.61 ab | 14935.33 ab | 16039.72 b |
| Urea×KCl150 | 1352.01 d | 155.33 d | 0.31 b | 1.46 c | 13705.49 d | 14488.36 d |
| Urea×KCl300 | 1414.21 c | 164.98 c | 0.34 ab | 1.52 bc | 14097.34 cd | 15111.23 c |
| Urea×KCl450 | 1367.03 d | 159.35 cd | 0.32 b | 1.49 c | 14039.76 cd | 15017.74 c |
| **Source of variance** |  |  |  |  |  |  |
| N | <0.0001 | <0.0001 | 0.0111 | <0.0001 | <0.0001 | <0.0001 |
| K | <0.0001 | 0.0013 | 0.0197 | 0.0003 | 0.0002 | 0.0004 |
| N×K | 0.063 | 0.3063 | 0.9734 | 0.1898 | 0.0321 | 0.0937 |
| **Note:** Means followed by different lowercase letters in the same column were significantly different based on analyses with ANOVAs followed by Duncan tests ($P < 0.05$). |  |  |  |  |  |  |

### Yield

The fresh and dry Italian ryegrass yields of the CRU treatment increased continuously in four clipping periods (Tables 7 and 8), those of the control treatment and common urea treatment decreased at the third clipping stage, and the difference was significant. In the whole growth period, under the CRU treatment conditions, there was no difference in the treatment with different amounts of KCl fertilizer, which indicated that the factors affecting the fresh and dry yields of Italian ryegrass were due more to nitrogen fertilizer than to potassium fertilizer. However, there was no significant N×K interaction effect on the yield. When clipping at the second stage, the highest dry yields of the CRU×KCl300 and CRU×KCl450 treatments were 1.75 g plant^−1^, which were 5.4% and 2.9% higher than those of the urea×KCl300 and urea×KCl450 treatments, respectively, at the same potassium rate. During the third and fourth stages, the growth trend of the plants increased slowly, the demand for nutrients decreased, and the CRU slowly released nitrogen for plant growth. Thus, CRU was more beneficial than common urea. In addition, potassium fertilizer had little influence on the yield. There was no significant N×K interaction effect on the dry yield and fresh yield (except at the fourth clipping).

**Table 7.**
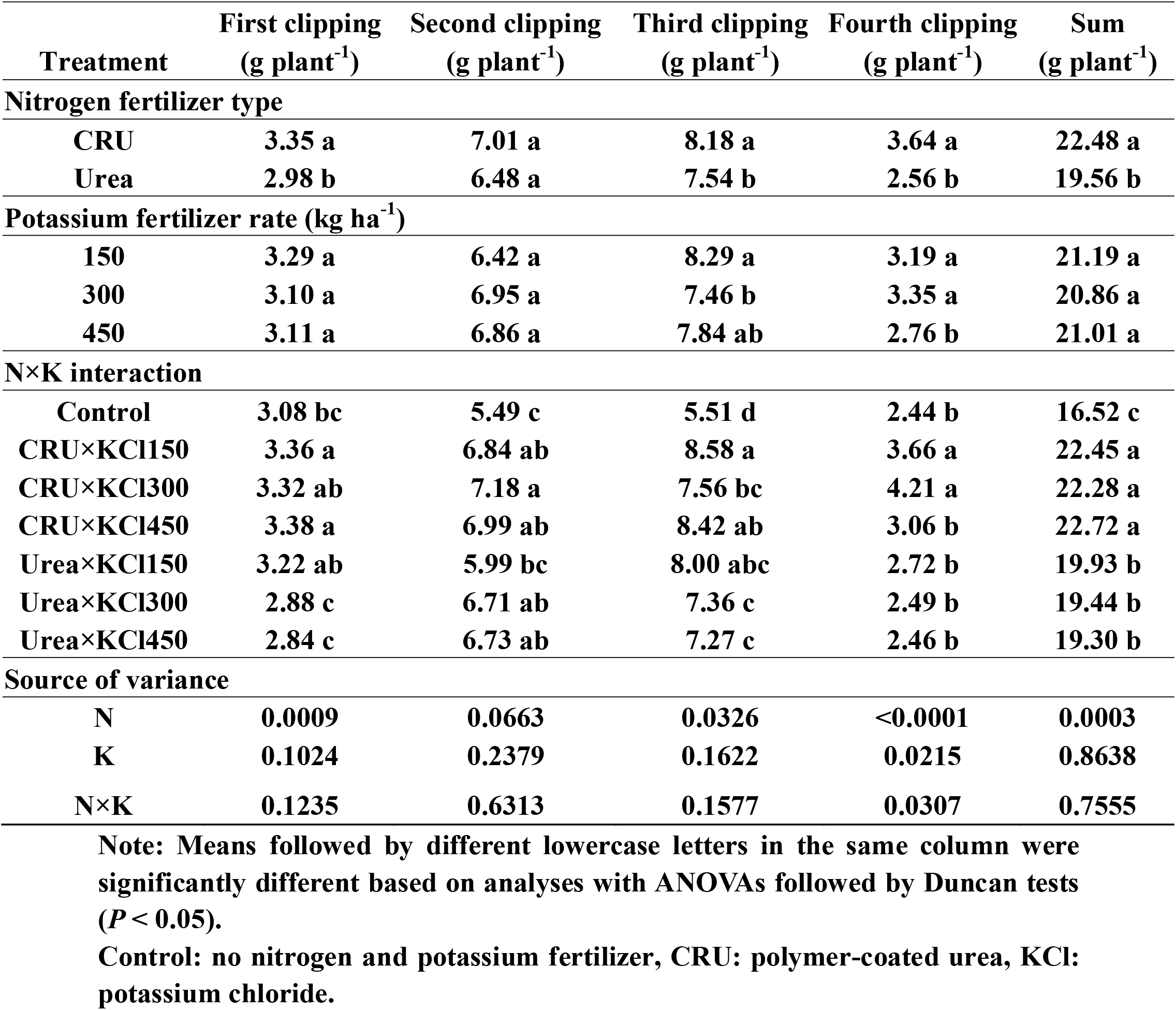
The fresh yield of Italian ryegrass under different treatments.

**Table 8.**
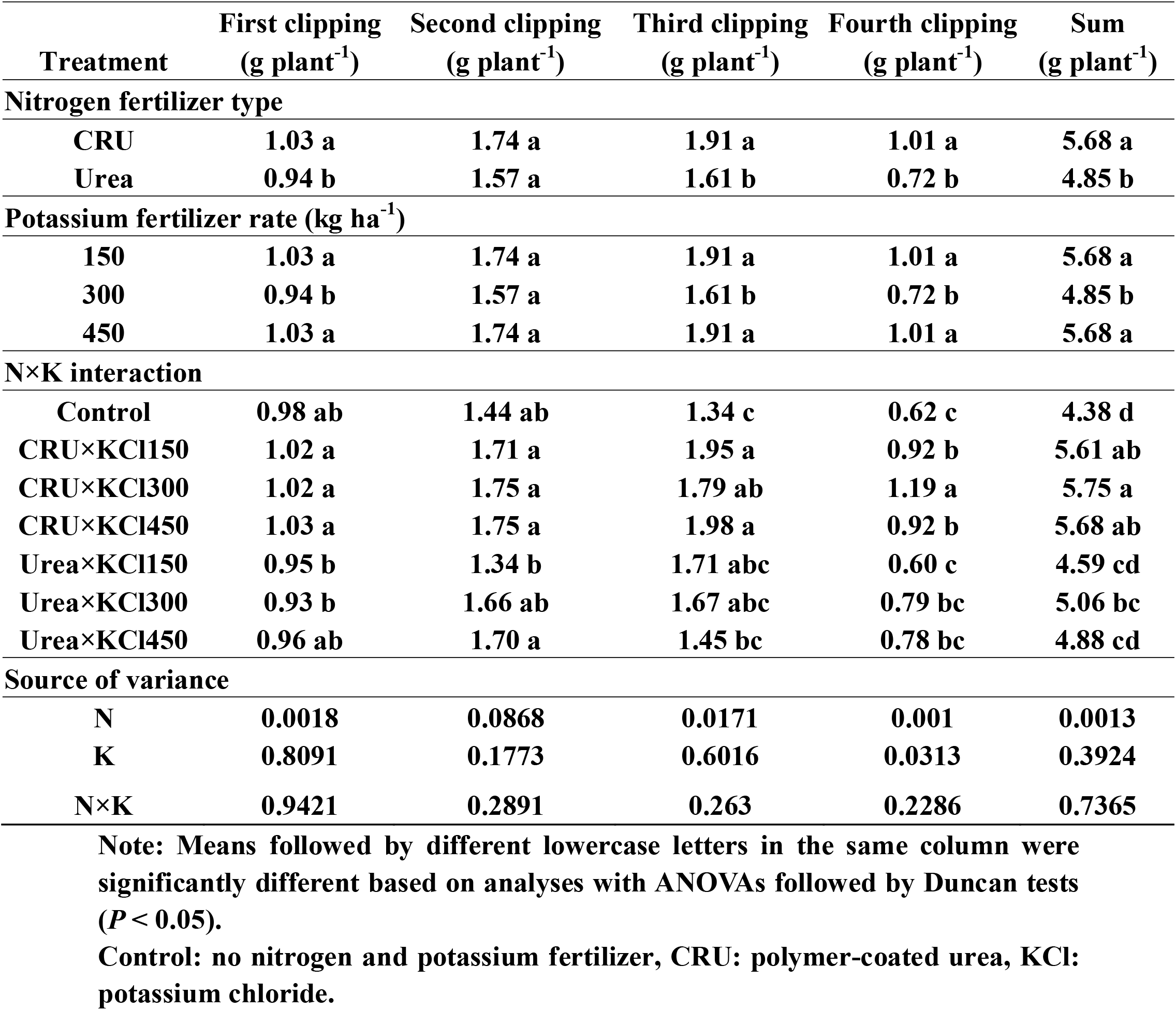
The dry yield of Italian ryegrass under different treatments.

| Treatment | First clipping<br>(g plant<sup>-1</sup>) | Second clipping<br>(g plant<sup>-1</sup>) | Third clipping<br>(g plant<sup>-1</sup>) | Fourth clipping<br>(g plant<sup>-1</sup>) | Sum<br>(g plant<sup>-1</sup>) |
| --- | --- | --- | --- | --- | --- |
| **Nitrogen fertilizer type** |  |  |  |  |  |
| CRU | 1.03 a | 1.74 a | 1.91 a | 1.01 a | 5.68 a |
| Urea | 0.94 b | 1.57 a | 1.61 b | 0.72 b | 4.85 b |
| **Potassium fertilizer rate (kg ha<sup>-1</sup>)** |  |  |  |  |  |
| 150 | 1.03 a | 1.74 a | 1.91 a | 1.01 a | 5.68 a |
| 300 | 0.94 b | 1.57 a | 1.61 b | 0.72 b | 4.85 b |
| 450 | 1.03 a | 1.74 a | 1.91 a | 1.01 a | 5.68 a |
| **N×K interaction** |  |  |  |  |  |
| Control | 0.98 ab | 1.44 ab | 1.34 c | 0.62 c | 4.38 d |
| CRU×KCl150 | 1.02 a | 1.71 a | 1.95 a | 0.92 b | 5.61 ab |
| CRU×KCl300 | 1.02 a | 1.75 a | 1.79 ab | 1.19 a | 5.75 a |
| CRU×KCl450 | 1.03 a | 1.75 a | 1.98 a | 0.92 b | 5.68 ab |
| Urea×KCl150 | 0.95 b | 1.34 b | 1.71 abc | 0.60 c | 4.59 cd |
| Urea×KCl300 | 0.93 b | 1.66 ab | 1.67 abc | 0.79 bc | 5.06 bc |
| Urea×KCl450 | 0.96 ab | 1.70 a | 1.45 bc | 0.78 bc | 4.88 cd |
| **Source of variance** |  |  |  |  |  |
| N | 0.0018 | 0.0868 | 0.0171 | 0.001 | 0.0013 |
| K | 0.8091 | 0.1773 | 0.6016 | 0.0313 | 0.3924 |
| N×K | 0.9421 | 0.2891 | 0.263 | 0.2286 | 0.7365 |

Note: Means followed by different lowercase letters in the same column were significantly different based on analyses with ANOVAs followed by Duncan tests ( $P < 0.05$ ).
Control: no nitrogen and potassium fertilizer, CRU: polymer-coated urea, KCl: potassium chloride.

### Nitrogen and potassium use efficiency

The nitrogen fertilizer application significantly affected nitrogen uptake and NUE, and the potassium fertilizer application significantly affected potassium uptake and KUE (Table 9). The nitrogen uptake and NUE of the CRU treatments were significantly higher than those of the urea treatments. In addition, potassium uptake increased with increasing KCl rate, but the KUE had the opposite trend. In total, the CRU×KCl300 treatment resulted in the highest nutrient uptake and use efficiency.

**Table 9.** The nitrogen uptake, potassium uptake, nitrogen use efficiency (NUE) and potassium use efficiency (KUE) of Italian ryegrass under different treatments.

| Treatment | N uptake<br>(kg ha<sup>-1</sup>) | K uptake<br>(kg ha<sup>-1</sup>) | NUE<br>(%) | KUE<br>(%) |
| --- | --- | --- | --- | --- |
| **Nitrogen fertilizer type** |  |  |  |  |
| CRU | 313.56 a | 298.78 a | 38.37 a | 31.77 a |
| Urea | 298.78 b | 297.22 a | 33.45 b | 30.46 a |
| **Potassium fertilizer rate (kg ha<sup>-1</sup>)** |  |  |  |  |
| 150 | 305.50 a | 270.67 c | 35.82 a | 33.58 a |
| 300 | 308.01 a | 304.00 b | 36.05 a | 34.79 a |
| 450 | 305.32 a | 319.33 a | 35.86 a | 24.98 b |
| **N×K interaction** |  |  |  |  |
| Control | 202.33 c | 212.61 d | - | - |
| **CRU×KCl150** | **312.00 a** | **270.04 c** | **38.15 b** | **33.69 a** |
| **CRU×KCl300** | **315.67 a** | **306.33 ab** | **38.59 a** | **36.59 a** |
| **CRU×KCl450** | **313.22 b** | **320.09 a** | **38.36 ab** | **25.03 b** |
| **Urea×KCl150** | **299.12 b** | **271.33 c** | **33.49 c** | **33.46 a** |
| **Urea×KCl300** | **300.36 b** | **301.67 b** | **33.50 c** | **32.99 a** |
| **Urea×KCl450** | **297.11 b** | **318.64 a** | **33.35 c** | **24.93 b** |
| **Source of variance** |  |  |  |  |
| **N** | **<0.0001** | **0.6245** | **<0.0001** | **0.2911** |
| **K** | **0.2975** | **<0.0001** | **0.1275** | **0.0002** |
| **N×K** | **0.7215** | **0.7333** | **0.1572** | **0.417** |

**Note:** Means followed by different lowercase letters in the same column were significantly different based on analyses with ANOVAs followed by Duncan tests ( $P < 0.05$ ).
**Control:** no nitrogen and potassium fertilizer, **CRU:** polymer-coated urea, **KCl:** potassium chloride.

## Discussion

### Soil inorganic nitrogen and potassium form

The contents of soil inorganic nitrogen (NO_3_^−^-N and NH_4_^+^-N) in the 0-20 cm soil layer were greatly affected by fertilization and were related to the type of fertilizer, the frequency of fertilization and the amount of fertilization (Zheng *et al*., 2016). In the present study, the contents of nitrate nitrogen and ammonium nitrogen at depths of 0-20 cm were significantly affected by nitrogen fertilization, but no significant difference was found between the potassium fertilization applications. In the beginning, urea rapidly dissolved and released substantial nitrogen, while the CRU initially released less nitrogen; thus, the soil inorganic nitrogen content of the urea treatment was higher than that of the CRU treatment at the first clipping stage. However, due to the continuous release of nitrogen from the CRU, the contents of NO_3_^−^-N and NH_4_^+^-N increased significantly from the second clipping stage to the fourth clipping stage compared with that in the urea treatment. Zheng *et al*. (2016) also found that the application of CRU mixed with urea significantly increased the soil inorganic nitrogen compared with the application of normal urea, especially in the later growth stage. The effect of the KCl rate on soil inorganic nitrogen was not obvious during the whole growth stage of Italian ryegrass.

The application of potassium fertilizer significantly affects the content of soil potassium and the form of soil potassium (Yang *et al*., 2017; Li *et al*., 2017). In the present study, potassium application significantly increased the contents of soil available K, water-soluble K, exchangeable K and non-exchangeable K, similar to other research results (Kurbah and Dixit, 2019). However, quick-acting fertilizers such as KCl are easily converted into non-exchangeable K, which leads to a decrease in soil potassium availability (Chen *et al*., 2020). In total, the contents of soil available K, water-soluble K, exchangeable K and non-exchangeable K increased with increasing KCl rate. The contents of soil available K and water-soluble K had a significant positive correlation, where a similar presence was also found in exchangeable K and non-exchangeable K (Table 10). The water-soluble K contributed the most to NO_3_^−^-N, and the available K had similar effects on NH_4_^+^-N. In addition, the contents of soil available potassium in the KCl300 and KCl450 treatments were similar. Thus, a moderate amount of potassium fertilizer could maintain a high potassium content in the soil, which would supply enough potassium nutrition for Italian ryegrass (Jiménez-Calderón *et al*., 2018). In addition, we found that the interaction effect of N×K on soil available potassium or inorganic nitrogen was not obvious during the whole growth stage of Italian ryegrass. Yang *et al*. (2016b) also found that the interaction effect of nitrogen and potassium fertilization on soil potassium and nitrogen was not obvious in a cotton field.

**Table 10.** Correlations between different forms of soil potassium and nitrogen.

| **Correlation coefficient** | **Available K** | **Water soluble K** | **Exchangeable K** | **Non-exchangeable K** | **NO<sub>3</sub><sup>-</sup>-N** | **NH<sub>4</sub><sup>+</sup>-N** |
| --- | --- | --- | --- | --- | --- | --- |
| **Available K** | **1** |  |  |  |  |  |
| **Water soluble K** | **0.8733\*\*** | **1** |  |  |  |  |
| **Exchangeable K** | **0.6775\*\*** | **0.2333\*** | **1** |  |  |  |
| **Non-exchangeable K** | **0.8455\*\*** | **0.8248\*\*** | **0.4423\*\*** | **1** |  |  |
| **NO<sub>3</sub><sup>-</sup>-N** | **0.7592\*\*** | **0.8370\*\*** | **0.2516\*** | **0.5092\*\*** | **1** |  |
| **NH<sub>4</sub><sup>+</sup>-N** | **0.7919\*\*** | **0.7695\*\*** | **0.4189\*\*** | **0.5579\*\*** | **0.8917\*\*** | **1** |

**Note:** \* $P < 0.05$ ; \*\* $P < 0.01$ .

### Growth index

To understand the growth of crops over time, it is necessary to determine their nutritional status. The traditional determination of leaf colour is to detect the chlorophyll content, which has high accuracy, but it takes considerable work and time (Croft *et al*., 2017). Plant height, chlorophyll, leaf area and photosynthetic index are important parameters to characterize crop photosynthetic production capacity, crop growth and nutritional status (Lin *et al*., 2019). The results showed that in comparison to the CRU treatment, the urea treatment significantly reduced the plant height, SPAD, leaf area and photosynthetic index, which might have led to plant senescence.

In addition, the application of KCl significantly increased the chlorophyll content in Italian ryegrass. Within a certain range of KCl applications, the chlorophyll content increased with increasing KCl applications, but beyond this threshold, the effect of KCl was weakened, thus affecting the ornamental value of Italian ryegrass. This outcome may have been due to the decrease in assimilative capacity, enzyme content and enzyme activity caused by the lack or excess of potassium, further leading to the decrease in leaf area and photosynthesis and thus affecting photosynthesis (Lu *et al*., 2017). Generally, the CRU×KCl300 treatment improved the leaf photosynthesis of Italian ryegrass.

Roots play an important role in crop growth and yield formation (Bandeoğlu *et al*., 2004). In the present study, the application of the CRU and moderate KCl rate significantly increased root length, surface areas and the numbers of tips and branches compared with other treatments, and there was a significant N×K interaction effect on the numbers of tips. These results suggested that suitable nitrogen and potassium fertilizer application can enhance root growth and thereby increase the uptake of nutrients (Enriquez-Hidalgo *et al*., 2018). In total, there was no significant N×K interaction effect on the growth index. Dong *et al*. (2010) also found a positive correlation between the duration of reproductive growth and the appropriate amount of N or K application, but the interaction effect of N×K was not obvious.

### Yield and fertilizer use efficiency

Nitrogen and potassium are the key factors affecting the yield of Italian ryegrass. Under sufficient nitrogen, the leaves of Italian ryegrass are thick green and have strong growth and a high yield of fresh grass, and under nitrogen deficiency, the leaves of Italian ryegrass are yellow and have poor growth (Vleugels *et al*., 2017).

The results showed that in comparison to those of the urea treatment and the control treatment, the fresh and dry yields of the CRU treatment were highest and increased by 20.5-53.2%. Studies have also shown that CRU effectively promotes the ability of photosynthesis to produce organic matter and then increase plant yield (Van Eerd *et al*., 2017; Miyatake *et al*., 2019). In this study, the second and third clipping stages were the period of high yield of Italian ryegrass and the period of high yield of CRU. This highlighted that the N uptake of Italian ryegrass in the first growing stages is lower, which can be functional to the presented pattern of N release in soil (Masoni *et al*., 2015). In the present study, the yield of the CRU treatment was higher than that of the common urea treatment in the four clipping periods, especially in the middle and later stages of Italian ryegrass growth, which occurred because the CRU slowly released nitrogen and met the growth demand of Italian ryegrass. In general, the nitrogen fertilizer type markedly affected the fresh and dry yield, but the application amount of potassium fertilizer had little effect on the yield. Moreover, no significant N×K interaction effect was found in the present study, which was different from Yang *et al*. (2016a). Hence, it is necessary and important to carry out further field experiments on the accurate rate and timing of N and K. Through a comparison, the yield of the CRU×KCl450 treatment was found to be the highest in the early stage, but considering the later stage yield, fertilizer utilization rate, economic benefits and other factors, the whole growth period of the CRU×KCl300 treatment was the most suitable treatment combination of all treatments, which improved the yield of Italian ryegrass.

There are many parameters to describe the efficiency of fertilizer utilization. The key to improving the efficiency of fertilizer utilization is nutrient absorption (Xue *et al*., 2017). In this study, NUE and KUE were used. Regardless of the amount of KCl applied, the NUE of the CRU treatment was significantly higher than that of the urea treatment, which might have been due to high nitrogen absorption, and Gaylord *et al*. (1975) also found similar results. In addition, the KCl rate had a significant effect on potassium absorption and KUE. The KUE values of the KCl300 treatment were higher than those of the KCl150 and KCl450 treatments. Therefore, the CRU×KCl300 treatment could improve the NUE and KUE of Italian ryegrass. In the present study, at the same N rate, the inorganic nitrogen concentration in the soil of the CRU was higher than that in the urea treatments, which indicated that more N remained in the soil, thus reducing the N leaching loss. Masoni *et al*. 2015 also found that the increase in nutrient use efficiency could also minimize unfavourable effects on the environment, mainly leaching.

## Conclusion

The nitrogen fertilizer type and potassium fertilizer rate had significant effects on Italian ryegrass growth, yield and soil fertility, but there was no significant N×K interaction effect. The CRU released nitrogen slowly, which was consistent with the nitrogen demand of Italian ryegrass during the whole growth and development period, simplifying the cultivation technology. This study found that the amount of potassium fertilizer had no significant effect on the growth of Italian ryegrass in the early stage, but in the middle and late stages of ryegrass growth, the CRU×KCl300 treatment improved plant SPAD and root morphology, delayed senescence of Italian ryegrass, and significantly increased the yield and fertilizer use efficiency. Hence, the CRU×KCl300 treatment is recommended as the best fertilization ratio for Italian ryegrass, and this recommendation can serve technical support for Italian ryegrass production and fertilization.

## Acknowledgment

The authors appreciate American Journal Experts (AJE) for English language editing.

## Author contributions

J.Q.C. and X.Y.Y. conceived and designed the experiments; Q.J.L. and X.Q.H. analyzed the data; J.B.G. and X.Y.Y. wrote the manuscript; S.T.L. and H.L. were involved in the related discussion; Y.L. helps to improve the quality of the manuscript. All authors reviewed the manuscript.

## References

Alves dos Santos J, da Fonseca AF, Barth G, Zardo Filho R. 2018. Silage maize quality in different uses of Italian ryegrass and soil management methods after liming. Arch. Agron. Soil Sci. 64(2), 173–184.

Ata-Ul-Karim ST, Liu X, Lu Z, Zheng H, Cao W, Zhu Y. 2017. Estimation of nitrogen fertilizer requirement for rice crop using critical nitrogen dilution curve. Field Crop Res. 201, 32–40.

Bandeolu E, Eyidoan F, Yücel M, Avni ktem H. 2004. Antioxidant responses of shoots and roots of lentil to nacl-salinity stress. Plant Growth Regul. 42(1), 69–77.

Binder S, Isbell F, Polasky S, Catford JA, Tilman D. 2018. Grassland biodiversity can pay. P. Nalt. Acad. Sci. USA. 115(15), 3876–3881.

Bolinder MA, T. Kätterer, O. Andrén, Ericson L, Kirchmann H. 2010. Long-term soil organic carbon and nitrogen dynamics in forage-based crop rotations in northern sweden (63-64°n). Agr. Ecosyst. Environ. 138(3-4), 335–342.

Cavalli D, Cabassi G, Borrelli L, Geromel G, Bechini L, Degano L, Gallina PM. 2016. Nitrogen fertilizer replacement value of undigested liquid cattle manure and digestates. Eur. J. Agron. 73, 34–41.

Chen J, Guo Z, Chen H, Yang X, Geng J. 2020. Effects of different potassium fertilizer types and dosages on cotton yield, soil available potassium and leaf photosynthesis. Arch. Agron. Soil Sci. (just-accepted).

Croft H, Chen JM, Luo X, Bartlett P, Chen B, Staebler RM. 2017. Leaf chlorophyll content as a proxy for leaf photosynthetic capacity. Global Change Biol. 23(9), 3513–3524.

Dong HZ, Kong XQ, Li WJ, Tang W, Zhang DM. 2010. Effects of plant density and nitrogen and potassium fertilization on cotton yield and uptake of major nutrients in two fields with varying fertility. Field Crops Res. 119 (1), 106–113.

Enriquez-Hidalgo D, Gilliland TJ, Egan M, Hennessy D. 2018. Production and quality benefits of white clover inclusion into ryegrass swards at different nitrogen fertilizer rates. J. Agr. Sci. 1–9.

Fan X, Kawamura K, Xuan TD, Yuba N, Lim J, Yoshitoshi R, Obitsu T. 2018. Low-cost visible and near-infrared camera on an unmanned aerial vehicle for assessing the herbage biomass and leaf area index in an Italian ryegrass field. Grass. Sci. 64(2), 145–150.

Gaylord MV. 1975. Spotted response of overseeded ryegrass to sulfur-coated urea. Agron. J. 67(6), 838–840.

Geng JB, Ma Q, Chen JQ, Zhang M, Li CL, Yang YC, Yang XY, Liu ZG. 2016. Effects of polymer coated urea and sulfur fertilization on yield, nitrogen use efficiency and leaf senescence of cotton. Field Crop Res. 187, 87–95.

Hasanuzzaman M, Bhuyan MHM, Nahar K, Hossain M, Mahmud JA, Hossen M, Fujita M. 2018. Potassium: a vital regulator of plant responses and tolerance to abiotic stresses. Agron. 8(3), 31.

He HB, Li WX, Zhang YW, Cheng JK, Jia XY, Li S, Xin GR. 2020. Effects of Italian ryegrass residues as green manure on soil properties and bacterial communities under an Italian ryegrass (*Lolium multiflorum* L.)-rice (*Oryza sativa* L.) rotation. Soil Till Res. 196, 104487.

Hussain F, Hussain I, Khan AHA, Muhammad YS, Iqbal M, Soja G, Yousaf S. 2018. Combined application of biochar, compost, and bacterial consortia with Italian ryegrass enhanced phytoremediation of petroleum hydrocarbon contaminated soil. Environ. Exp. Bot. 153, 80–88.

Jiménez-Calderón JD, Martínez-Fernández A, Benaouda M, Vicente F. 2018. A winter intercrop of faba bean and rapeseed for silage as a substitute for Italian ryegrass in rotation with maize. Arch. Agron. Soil Sci. 64(7), 983–993.

Lin KH, Wu CW, Chang YS. 2019. Applying dickson quality index, chlorophyll fluorescence, and leaf area index for assessing plant quality of Pentas lanceolata. Not. Bot. Horti Agrobo. 47(1), 169–176.

Jarvis SC. 1987. The effects of low, regulated supplies of nitrate and ammonium nitrogen on the growth and composition of perennial ryegrass. Plant Soil. 100(1), 99–112.

Kurbah I, Dixit SP. 2019. Soil potassium fractions as influenced by integrated fertilizer application based on soil test crop response under maize-wheat cropping systems in Acid Alfisol. Int. J. Econ. Plant. 6(1), 25–29.

Li P, Lu J, Wang Y, Wang S, Hussain S, Ren T, Li X. 2018. Nitrogen losses, use efficiency, and productivity of early rice under controlled-release urea. Agr. Ecosyst. Environ. 251, 78–87.

Li J, Niu L, Zhang Q, Di H, Hao J. 2017. Impacts of long-term lack of potassium fertilization on different forms of soil potassium and crop yields on the North China Plains. J. Soil Sediment. 17(6), 1607–1617.

Liu J, Yang Y, Gao B, Li YC, Xie J. 2019. Bio-based elastic polyurethane for controlled-release urea fertilizer: Fabrication, properties, swelling and nitrogen release characteristics. J. Clean. Prod. 209, 528–537.

Liu M, Dries L, Heijman W, Huang J, Zhu X, Hu Y, Chen H. 2018. The impact of ecological construction programs on grassland conservation in Inner Mongolia, China. Land Degrad. Dev. 29(2), 326–336.

Lu Z, Pan Y, Hu W, Cong R, Ren T, Guo S, Lu J. 2017. The photosynthetic and structural differences between leaves and siliques of Brassica napus exposed to potassium deficiency. BMC Plant Biol. 17(1), 240.

Martin K, Edwards G, Bryant R, Hodge M, Moir J, Chapman D, Cameron K. 2017. Herbage dry-matter yield and nitrogen concentration of grass, legume and herb species grown at different nitrogen-fertiliser rates under irrigation. Anim. Prod. Sci. 57(7), 1283–1288.

Masoni A, Mariotti M, Ercoli L, Pampana S, Arduini I. 2015. Nitrate leaching from forage legume crops and residual effect on italian ryegrass. Agrochimica Pisa. 59(1), 75–91.

McDonnell RP, Staines MV, Bolland MDA. 2018. Determining the critical plant test potassium concentration for annual and Italian ryegrass on dairy pastures in south-western Australia. Grass Forage Sci. 73(1), 112–122.

Miyatake M, Ohyama T, Yokoyama T, Sugihara S, Motobayashi T, Kamiya T, Ohkama-Ohtsu N. (2019). Effects of deep placement of controlled-release nitrogen fertilizer on soybean growth and yield under sulfur deficiency. Soil Sci. Plant Nutr. 65(3), 259–266.

Oliveira EMD, Oliveira Filho JDC, Oliveira RAD, Oliveira RMD, Cecon PR. 2017. Determination of xaraés grass quality submitted to irrigation water levels and nitrogen and potassium doses. Eng. Agr-Jaboticabal. 7(1), 64–74.

Rietra RP, Heinen M, Dimkpa CO, Bindraban PS. 2017. Effects of nutrient antagonism and synergism on yield and fertilizer use efficiency. Commun. Soil Sci. Plan. 48(16), 1895–1920.

Soil Survey Staff. 1999. Soil taxonomy. In: Soil Survey Staff (Ed.), a basic system of soil classification for making and interpreting Soil Surveys, 2nd U.S. Gov. Print. Office, Washington, DC. pp. 163–167.

Svatos KB, Abbott LK. 2019. Dairy soil bacterial responses to nitrogen application in simulated Italian ryegrass and white clover pasture. J. Dairy Sci. 102(10), 9495–9504.

Snyder GH, Cisar JL. 2000. Nitrogen/potassium fertilization ratios for bermudagrass turf. Crop Sci. 40(6), 1719–1723.

Tang Q, Feng M. 2002. Practical Statistics and DPS Data Processing System. China Agric. Press. Beijing.

Van Eerd LL, Turnbull JJD, Bakker CJ, Vyn RJ, McKeown AW, Westerveld SM. 2017. Comparing soluble to controlled-release nitrogen fertilizers: storage cabbage yield, profit margins, and N use efficiency. Can. J. Plant Sci. 98(4), 815–829.

Vleugels T, Rijckaert G, Gislum R. 2017. Seed yield response to N fertilization and potential of proximal sensing in Italian ryegrass seed crops. Field Crop Res. 211, 37–47.

Wang H, Wu L, Cheng M, Fan J, Zhang F, Zou Y, Wang X. 2018. Coupling effects of water and fertilizer on yield, water and fertilizer use efficiency of drip-fertigated cotton in northern Xinjiang, China. Field Crop Res. 219, 169–179.

Woods RR, Cameron KC, Edwards GR, Di HJ, Clough TJ. 2018. Reducing nitrogen leaching losses in grazed dairy systems using an Italian ryegrass-plantain-white clover forage mix. Grass Forage Sci. 73(4), 878–887.

Xue X, Mai W, Zhao Z, Zhang K, Tian C. 2017. Optimized nitrogen fertilizer application enhances absorption of soil nitrogen and yield of castor with drip irrigation under mulch film. Ind. Crop Prod. 95, 156–162.

Yang XY, Geng JB, Li CL, Zhang M, Chen BC, Tian XF, Zheng WK, Liu ZG, Wang C. 2016a. Combined application of polymer coated potassium chloride and urea improved fertilizer use efficiencies, yield and leaf photosynthesis of cotton on saline soil. Field Crops Res. 197, 63–73.

Yang XY, Geng JB, Li CL, Zhang M, Tian XF. 2016b. Cumulative release characteristics of controlled-release nitrogen and potassium fertilizers and their effects on soil fertility, and cotton growth. Sci. Rep. 6, 39030.

Yang XY, Li CL, Zhang Q, Liu ZG, Geng JB, Zhang M. 2017. Effects of polymer-coated potassium chloride on cotton yield, leaf senescence and soil potassium. Field Crops Res. 212, 145–152.

Zheng W, Sui C, Liu Z, Geng J, Tian X, Yang X, Zhang M. 2016. Long-term effects of controlled-release urea on crop yields and soil fertility under wheat–corn double cropping systems. Agron. J. 108(4), 1703–1716.

